# Multimodal laminar characterization of visual areas along the cortical hierarchy

**DOI:** 10.1101/2024.11.18.624072

**Authors:** Alessandra Pizzuti, Pierre-Louis Bazin, Dimo Ivanov, Sebastian Dresbach, Judith Peters, Rainer Goebel, Omer Faruk Gulban

## Abstract

Understanding the relationship between brain structure and function is a central goal in neuroscience. While post-mortem studies using microscopic techniques have provided detailed insights into the brain’s cytoarchitectonic and myeloarchitectonic patterns, linking these structural findings to functional outcomes remains challenging. Magnetic resonance imaging (MRI) has emerged as a powerful non-invasive tool for studying both structure and function, but discrepancies in spatial resolution between structural and functional imaging, especially in layer-fMRI, complicate the interpretation of functional results. In this study, we explore how visual cortical hierarchy relates to microscopic and mesoscopic laminar features. Focusing on visual areas that span progressive hierarchical levels, V1, V2, V3, and hMT+, we apply a multimodal approach combining post-mortem histology, post-mortem and in-vivo quantitative MRI (qMRI), and resting-state layer-fMRI. Using the open-access post-mortem AHEAD dataset, which integrates histological and qMRI contrasts from the same brain samples, we bridge microscopic observations with qMRI data. In parallel, we incorporate high-resolution 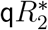 MRI and resting-state layer-fMRI from the same participant, allowing for a comparative analysis of laminar profiles across cortical depth. For computing laminar profiles, we developed an analysis pipeline that bridges histology images, mesoscopic qMRI, and layer-fMRI. Our findings highlight parvalbumin laminar profiles (reflecting interneuron parvalbumin density) as the most discriminative feature for differentiating brain areas. Additionally, we report laminar quantitative 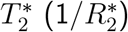 profiles from post-mortem and in-vivo data, together with 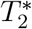-weighted resting-state layer-fMRI, all of which exhibit a similar overall shape across modalities. Using our methodological framework, a similar laminar characterization can be extended to study other brain regions. Generative models for layer fMRI will benefit from incorporating these new empirical microstructural (parvalbumin) and physical quantitative 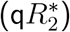 data, leading to more area-specific and accurate models.

**Highlights:**

1. We present a multimodal analysis of the laminar organization of four visual regions (V1, V2, V3, hMT+), characterizing progressive visual hierarchy levels in humans. This analysis spans from post-mortem microscopy and quantitative MRI (qMRI) to in-vivo qMRI and laminar fMRI during resting state.
2. Among the three microscopy contrasts, parvalbumin, a marker of interneuron density, emerges as the most distinctive regional feature. Notably, the parvalbumin laminar profiles vary across hierarchy levels, with hMT+ showing the greatest divergence compared to V1, V2, and V3.
3. Quantitative 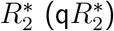 measurements, from both post-mortem and in-vivo data, reveal a clear increase towards the superficial cortical layers. These depth-dependent patterns closely mirror the laminar profiles observed in both task and resting-state fMRI.
4. Surprisingly, no substantial difference was observed in laminar 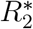 profiles between post-mortem and in-vivo data across the visual areas.

## 1 Introduction

Understanding the functionality of any physical component fundamentally requires exploring the relationship between its structure and function. This principle is a cornerstone of neuroscience, where one of the field’s enduring goals is to elucidate how brain structure underpins its myriad of functions. Since the early 20th century, scientists have conducted extensive post-mortem studies on the brain’s structure, particularly fusing microscopic techniques that offer spatial resolution at the micrometer scale. These studies have led to the discovery that the brain’s cortical architecture is not uniform. Different brain regions exhibit distinct laminar cytoarchitectonic and myeloarchitectonic patterns, allowing for the establishment of cortical parcellations over time (Brodmann, 1909; Nieuwenhuys, 2013; Vogt, 1906; Zilles and Amunts, 2010). Despite the valuable insights gained from these studies, post-mortem approaches have inherent limitations in capturing functional coupling of microstructures, as functional testing is typically not conducted on the same data sample. Although recent studies have demonstrated the feasibility of combining in-vivo and ex-vivo data from the same participants (Boon et al., 2019; Jonkman et al., 2019), this approach remains challenging and is not yet widely adopted (Fischl and Sereno, 2018).

One candidate technique to complement post-mortem brain imaging is magnetic resonance imaging (MRI) as a non-invasive method for studying both brain structure and function (Bandettini et al., 1992; Ogawa et al., 1992). Recently, advancements in ultra high magnetic fields (Koopmans and Yacoub, 2019; Shmuel et al., 2007; Uğurbil et al., 2003; Uludağ et al., 2009) have enabled significant improvements in spatial resolution, allowing for the imaging of the cortical landscape at the mesoscopic scale (*<* 1 mm). While in-vivo structural imaging could reach a spatial resolution of *<*=0.35 iso. mm (Bollmann et al., 2022; Gulban et al., 2022; Lüsebrink et al., 2021), the upcoming ‘layer-fmri field’ for functional imaging has reached submillimeter resolution and aims to test functional hypotheses related to areal microcircuits in vivo (De Martino et al., 2018; Dresbach et al., 2024a; Huber et al., 2017; Petro and Muckli, 2017; Pizzuti et al., 2023; Viessmann and Polimeni, 2021). Despite the push for a higher spatial resolution (below 0.5 iso. mm) (Feinberg et al., 2023; Vizioli et al., 2021), currently, the routinely used functional resolution is 0.8 mm isotropic. This discrepancy in resolution between structural and functional scans, leads to new challenges in interpreting the current layer-fMRI results. A potential solution to these challenges is the integration of multiple modalities. For the question of brain parcellation, combining microstructural post-mortem data with topographical and functional in-vivo datasets has resulted in the development of a new multimodal parcellation of the human cerebral cortex (Glasser et al., 2016). This approach has become the consensus standard for fMRI studies. Similarly, quantitative MRI (qMRI) techniques, combined with modeling approaches, provide critical microstructural information to complement in-vivo imaging (Dinse et al., 2015; Trampel et al., 2019; Weiskopf et al., 2021). Notably, among qMRI data, 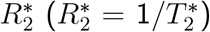 relaxation rate serves as a key measurement in understanding tissue microstructure, reflecting changes in magnetic susceptibility that can inform us about iron and myelin deposition. This is particularly relevant in fMRI, where the BOLD signal is predominantly 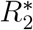 weighted, making 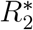 a key bridge between microstructural properties and functional imaging results. Here, we propose a multimodal study that aims to explore how cortical visual hierarchy relates to microscopic and mesoscopic laminar features. We provide a comprehensive laminar characterization of four visual areas that span progressive hierarchical levels such as V1, V2, V3 and hMT+. We combined post-mortem histological and qMRI data with in-vivo high-resolution quantitative and resting-state layer-fMRI (rs-fMRI). For the first part, we used the recently published open access post-mortem AHEAD dataset (Alkemade et al., 2022). This dataset includes multiple histological contrasts and three qMRI contrasts from the same brain, providing a unique opportunity to bridge MRI with direct microscopic observations. For the second part, we complement this post-mortem dataset with our in-vivo dataset combining high-resolution 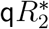 MRI at 0.35 iso. Mm resolution with resting-state layer-fMRI at 0.8 iso. mm from the same participant. By comparing laminar profiles from the two datasets, we provide a unified perspective on the laminar organization of these visual areas. Among the three microscopy contrasts, parvalbumin (marker of interneurons density) emerges as the most distinctive regional feature that varies across hierarchical levels. Notably, hMT+ parvalbumin laminar profile mostly differs from V1, V2, and V3. Moreover, we provide quantitative laminar profiles of 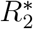 for both post-mortem and in-vivo brain samples. Our laminar quantification on microstructural composition and physical 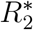 can be integrated in generative laminar fMRI models (Havlicek and Uludağ, 2020) to improve the areal laminar predictions and the interpretability of layer-fMRI results. Finally, we developed, streamlined and shared all the analysis methods used in this paper to cover the three modalities (histology, qMRI, layer-fMRI) together with our MRI/fMRI dataset. This framework of analysis offers a versatile approach that can be extended to investigate different brain areas, providing the tools to enhance our understanding of structure to function coupling across the entire cortex.

## 2 Materials and methods

### 2.1 Data overview

Our data set includes two main sources: the post-mortem 3D whole brain microscopy and 7T quantitative MRI (qMRI) from AHEAD dataset and an in-vivo whole-brain qMRI 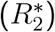 at 0.35 iso. mm resolution and resting state fMRI at 0.8 iso. mm from the same participant. A data overview is illustrated in (**Figure 1**).

**Figure 1.**
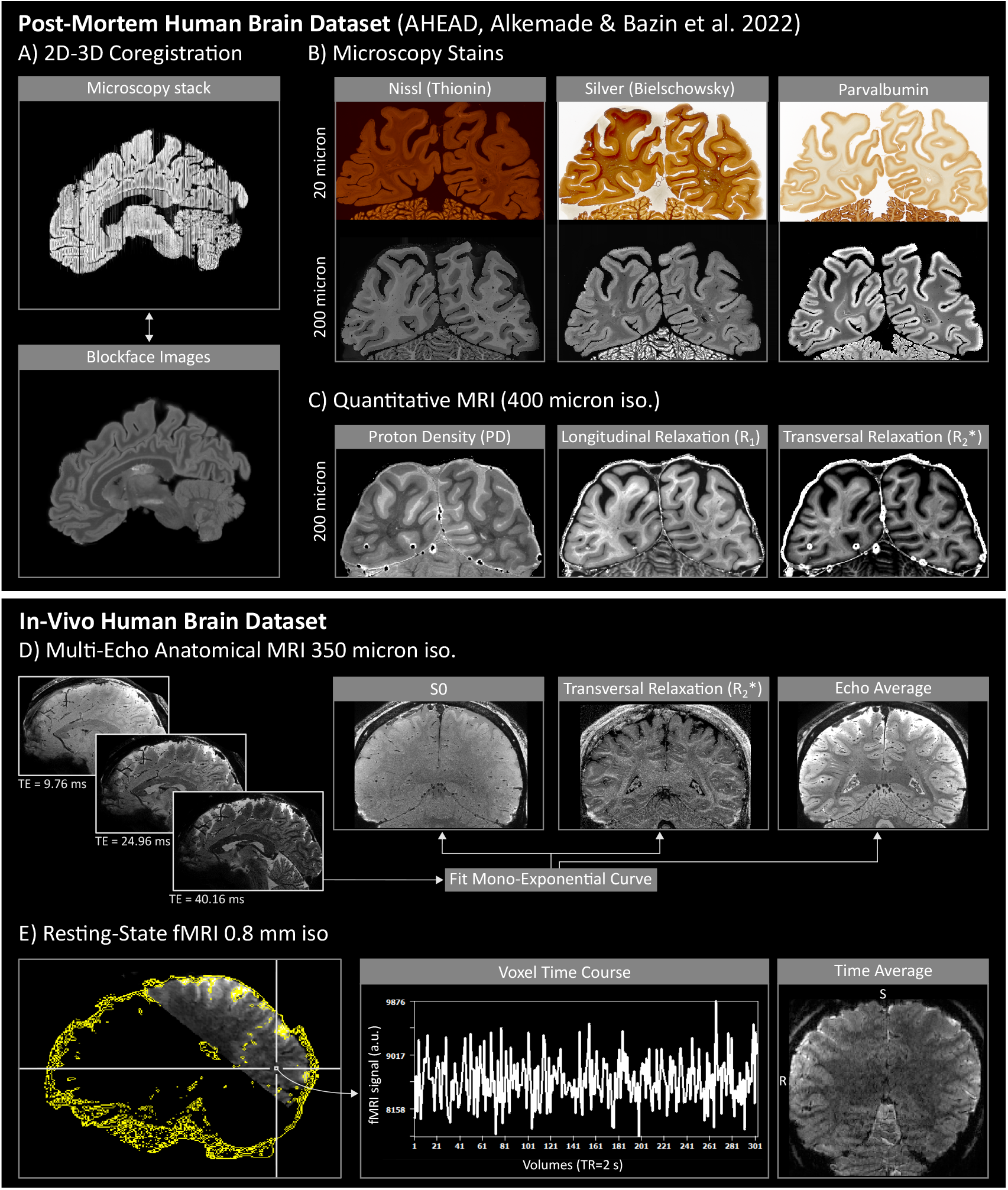
Data overview. A-C) Post-mortem human brain AHEAD dataset consisting of three microscopy stains: Nissl (Thionin) as neural density, Silver (Bielschowsky) as fiber density and Parvalbumin as interneurons density correlate (B) and three qMRI contrasts: proton density (PD), longitudinal relaxation (*R*_1_) and transversal relaxation 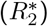 (C). Original stains were collected at 20 micrometers, while qMRI were collected at 400 micrometers. Both modalities were co-registered in the same brain space at 150 × 150 × 200 micrometer resolution through 2D-3D registration techniques with respect to the blockface images (A). In-vivo human brain dataset consisting of four runs of whole brain multi-echo multi-shot anatomical MRI at 350 micrometer isotropic resolution (D) and resting-state fMRI data at 0.8 mm isotropic with occipital coverage (E). Three echoes were collected and used to fit a mono-exponential curve and compute S0 and 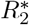 (D). rs-fMRI brain coverage and a voxel time course is shown in panel (E). Co-registration between anatomical and fMRI data was performed between the average echoes and average time course images.

The post-mortem AHEAD dataset is the first publicly available dataset containing a 3D whole-brain map of multiple microscopy contrasts and 7T qMRI from two human specimens. Whole-brain MRI acquired before sectioning consists of proton density, *R*_1_ and 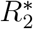 MRI maps with an isotropic resolution of 0.4 mm (**Figure 1C**). Coronal slices with in plane resolution of 0.021 mm and 0.20 mm across slices were stained for five microscopy contrasts in an interleaved fashion: two histology stains - Nissl (Thionin, glial and neuronal cell body density) and silver stain (Bielschowsky, fiber density) - and three immunochemistry stains -calbindin, calretinin, parvalbumin (interneurons density). Example slices at original resolution are shown in (**Figure 1B**). Note that calbindin and calretinin were only available at the medial part (central) of the brain, not covering the visual areas of interest. Notably, the authors used advanced post-processing techniques to align the coronal slices and reconstruct a 3D multi-contrast staining map aligned to the qMRI maps. As a result, the microscopic underpinning of MRI slices can be studied through the direct link between microscopy and MRI data. For a comprehensive and detailed description of post-mortem data collection and reconstruction of AHEAD dataset please refer to the original paper from (Alkemade et al., 2022). The in-vivo whole brain anatomical 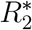 images and resting-state BOLD fMRI data were collected from the same participant with the whole-body MAGNETOM 7T “Plus” (Siemens Healthineers, Erlangen, Germany) at Scannexus B.V. (Maastricht, The Netherlands) using a 32-channel RX head-coil (Nova Medical, Wilmington, MA, USA). The shimming procedure included the vendor-provided routines to maximize the field homogeneity within the imaging slab. Anatomical images were collected at 0.35 iso mm resolution using a newly developed multi-echo multi-shot gradient recalled echo sequence with 3D echo planar imaging (EPI) readout (Gulban, 2024) (**Figure 1D**). By combining the strength of a 3D acquisition and highly segmented k-space through a multi-shot technique, we could collect 0.35 iso. mm whole-brain anatomical images with very limited geometric distortions in less than 10 min acquisition time (6 min 48 s). We used a multi-echo 3D EPI sequence (3 echoes), hereby referred to as 3D ME EPI, with the following main parameters: field of view (FoV) = 200×200×130 mm; orientation = sagittal; bandwidth = 546 Hz/Px; repetition time (TR) = 52.8 ms; vol.TR = 396 s; echo time (TE) = [9.76, 24.96, 40.16] ms; flip angle (FA) = 10°; Dual polarity = on; phase partial fourier = off; Segmentation = 40; EPI factor = 5; PAT mode=CAIPIRINHA; Acceleration factor phase encoding (PE)x3D=3×2; CAIPI trajectory = w/o z-blips. The complete protocol and the data used in this manuscript are publicly available in Zenodo: https://doi.org/10.5281/zenodo.14147820. We collected a total of four runs. We also used an available MP2RAGE at 0.35 mm isotropic previously collected in a separate session as “slab-stitched MP2RAGE” (Gulban, 2024). Briefly, five partial brain slabs at 0.35 iso. mm were concatenated to achieve whole-brain coverage. Each slab was collected within a single 10 minute run. The slabs were stitched in a post-processing step to have whole-brain images. Resting-state fMRI data (rs-fMRI) were collected by using a 2D GE EPI sequence with blood oxygen level dependent (BOLD) contrast (based on (Moeller et al., 2010)) with 0.8 isotropic mm resolution and coverage of the visual areas of interest (**Figure 1E**). The in-plane field of view was 140 × 137 mm (176 × 172 matrix) for a total of 58 acquired slices. The imaging parameters were: TE = 24.6 ms, TR = 2000 ms, flip angle FA = 69°, in plane partial Fourier factor 6/8, GRAPPA=3, multi-band (MB)=2. The raw data and the scanning protocol are available in Zenodo: https://doi.org/10.5281/zenodo.14164885. We placed a small functional imaging slab according to a predetermined positioning based on results from a functional visual localizer obtained in an independent experimental session (Pizzuti et al., 2024) using the auto-align sequence (AAscout) from Siemens. Before the acquisition of the main run, we collected 10 volumes for distortion correction with the settings specified above but opposite phase encoding direction (posterior-anterior). A total of 300 volumes were collected in 10 minutes while the participant was asked to fixate a black fixation cross on a gray background. A frosted screen (distance from eye to screen: 99 cm; image width: 28 cm; image height: 17.5 cm) at the rear of the magnet was used to project the visual stimuli (fixation cross) (using Panasonic projector 28 PT-EZ570; Newark, NJ, USA; resolution 1920×1200; nominal refresh rate: 60 Hz) that participants can watch through a tilted mirror attached to the head coil.

### 2.2 Region of interest definition and cortical segmentation

We focused our laminar analysis on the following visual areas: V1, V2, V3, and hMT+. The same Region-of-Interest (ROI) definition procedure was applied to both post-mortem and in-vivo dataset. For early visual areas (V1, V2, V3) we coregistered our data to a visual probabilistic functional atlas (visfatlas) (Rosenke et al., 2021) using cortex-based alignment as implemented in BrainVoyager (Goebel, 2012). For hMT+ we used the atlas published by Huang et al., 2019 since hMT+ was only partially included in the visfatlas (one hemisphere was missing in the current release). A schematic illustration of our ROI definition is reported in **Figure 2A** and in **Supplementary Figure 2**. By using visfatlas, we aimed to include the extent of the visual ROIs that can feasibly be stimulated during an fMRI, due to the reduced visual field that can be presented as stimulus in the scanner. For the AHEAD dataset, we used the ‘blockface’ images to which both microscopy and qMRI maps were aligned to (**Figure 1A**). Blockface images were obtained during sectioning and reconstructed as 3D volume at 0.15 × 0.15 × 0.2 mm resolution. In order to correctly import the post-mortem blockface images into BrainVoyager, we first make isotropic voxels (0.2 iso. mm) and then inverted the contrast. Within BrainVoyager, we downsampled the spatial resolution to 0.5 iso. mm and aligned to the ACPC space. For the in-vivo dataset, we used UNI contrast images from MP2RAGE at original resolution of 0.35 iso. mm coregisted to 3D ME EPI and to fMRI data. We followed Brainvoyager’s ‘Advanced Segmentation Pipeline’ to compute white and gray matter tissue segmentation. A manual refinement of the white matter segmentation was performed by the author A.P. in order to remove the mislabeled voxels creating holes or false geometries, especially around the occipital pole (e.g. around the sinus, subcortical areas). The resulting segmentation files (one from blockfase images and the one from MP2RAGE images) are then used to reconstruct the white matter surfaces within BrainVoyager. On each surface, we defined hMT+ by manual drawing and by matching the characteristic macro-anatomical features reported by Huang and colleagues (e.g. cortical localization, cortical surface areas) (**Figure 2A, Supplementary Figure 2**). The white matter surface was used as input for the cortex-based alignment pipeline of BrainVoyager. Once the alignment to the visfatlas was successfully performed, the ROIs on the cortical surface were all mapped back into the volume space (depth sampling -1 to +3 mm) within BrainVoyager and exported as NIFTI. We projected back into the original resolution 0.15 × 0.15 × 0.2 mm (for AHEAD dataset) and 0.175 iso. mm (for in-vivo dataset) using the program -greedy with ‘LABEL’ interpolation option to preserve the binary nature of the data (Tustison et al., 2010). The final resolution for in-vivo data (0.175 iso. mm) was chosen to match the resolution of computed 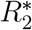 map (see In vivo MRI and fMRI signal extraction paragraph). Finally, a careful manual tissue segmentation around ROIs in the same native space was performed by A.P. and independently reviewed by O.F.G by using ITK-SNAP software (Tustison et al., 2010). Manual edits were lastly polished by using `LN2_RIM_POLISH` from LayNii (Huber et al., 2021) that implements a smoothing procedure using a combination of morphological operations of dilation and erosion. An exemplary slice showing the quality of our segmentation is reported in **Figure 2B, ii**. Within the final gray matter space, we projected the ROIs using LN2 VORONOI from LayNii.

**Figure 2.**
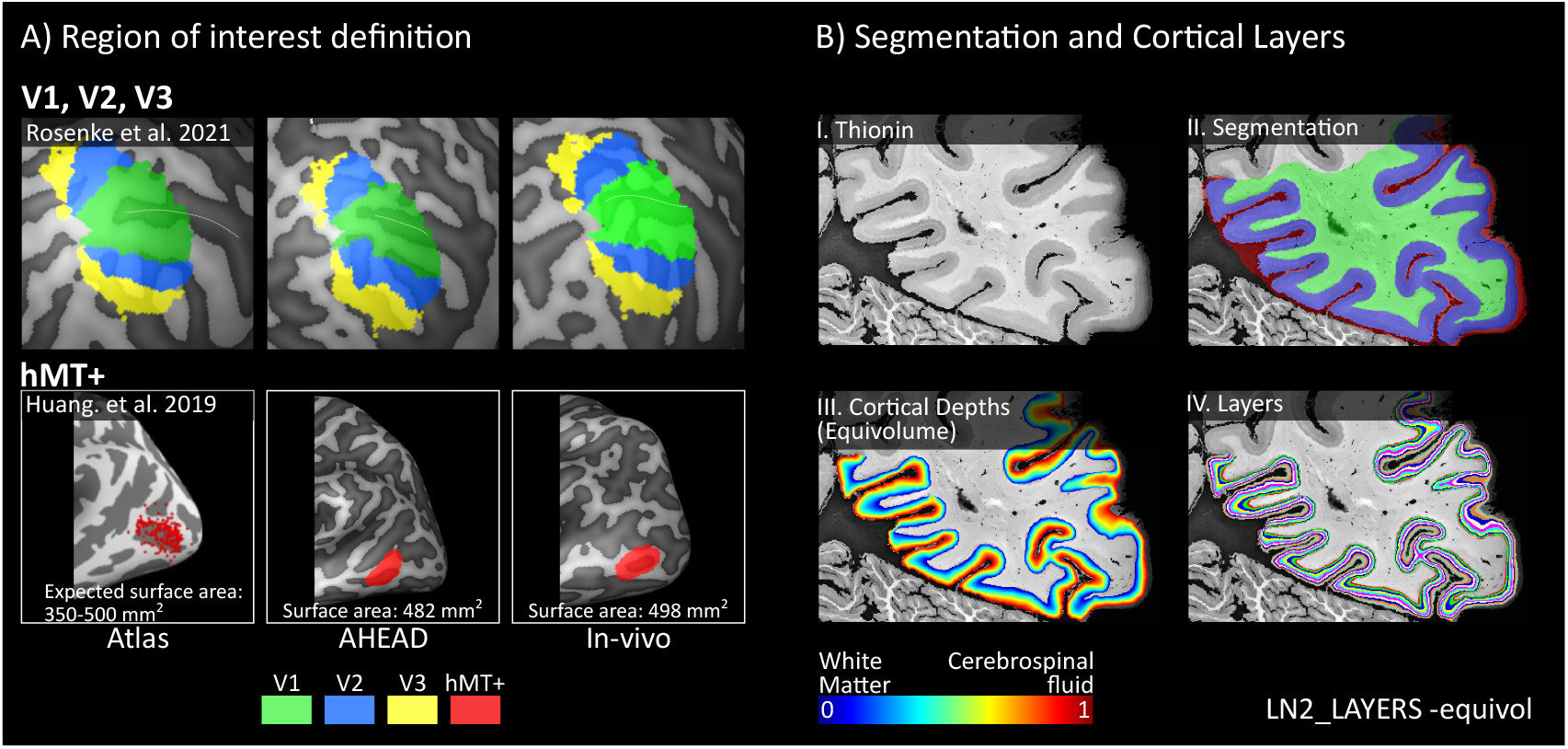
Overview of the two methodological steps. Panel (A) schematically illustrates the procedure used to define the regions of interest. A cortex-based alignment to the functional visual atlas (visfatlas, Rosenke et al. 2021) was used to define V1, V2, V3, while a macroanatomical procedure guided the definition of hMT+ to match a probabilistic atlas (Huang et al. 2019). Please, inspect **Supplementary Figure 2** for an extended overview of our ROI definition. Panel (B) illustrates an example of input (i) for the computation of the geometric cortical layers using the program `LN2_LAYERS` with the -equivol option. As output, normalized equivolume cortical depth measure (iii) is discretized in 11 equivolume layers (iv). Note that 11 layers are chosen here as an example for visualizing discrete layers; for the main results, the entire cortical depth (iii) is used to compute layer profile sas 2D histograms.

### 2.3 Geometric cortical layers

The definition of cortical depth measures for both post-mortem (at 0.2 iso. mm, nominal) and in-vivo dataset (0.175 iso. mm, nominal) is based on the accurate tissue segmentation (**Figure 2, B (ii)**) and was performed within LayNii using the LN2 LAYERS -equivol program. This program attributes a normalized (0-1) cortical depth measure to each voxel in 3D according to the equivolume principle (**Figure 2, B (iii)**). Only in the post-mortem data, we ran the algorithm iteratively on each 2D slice (0.15 × 0.15 mm), since the 3D reconstruction inevitably comes with misalignments and geometrical deformation. We are aware that the equivolume principle is defined for 3D data, however we qualitatively assessed that the amount of errors rising by using a slice-by-slice approach was less compared by using the 3D reconstructed data. Note that, most of the computational mistakes from the slice-by-slice layering procedure were automatically excluded from further analysis due to the ‘cutting angle’ filtering approach (see AHEAD laminar signal extraction paragraph). Laminar profiles were computed and shown as 2D histograms. The number of bins along the two dimensions (x-as cortical depth and y-as voxel intensity) was adjusted according to the region and the contrast represented. For each 2D histogram, we binned the cortical depth with 21 equivolume layers and overlaid a ‘median’ laminar profile by computing the median intensity values across voxels belonging to each layer separately. Median was chosen over mean as it is more resilient to outliers that mostly affect tissue boundaries.

### 2.4 AHEAD volumetric cortical parametrization

The tissue segmentation (**Figure 2, B (ii)**) is also used to define a 3D geodesic coordinate system for each cortical region of interest. This parametrization is needed to run our novel geodesic filters (see Tears filter and Bias Field filter described in AHEAD laminar signal extraction paragraph) for extracting laminar finescale details in AHEAD data. We obtained the first two sets of geodesic coordinates (U,V coordinates) for the gray matter of ROI separately, by running `LN2_MULTILATERATE` (input: segmentation file). The third coordinate (coordinate D), parametrizing the cortical depth dimension, was computed by running LN2 LAYERS -equivol for each slice of each ROI (input: segmentation file). While we ran the first program using the ROI segmentation computed on the 3D reconstructed model (see ‘ROI definition and cortical segmentation’ paragraph), the second program was run iteratively for each 2D slice using the final segmentation refined for each slice. In order to preserve the two-dimensional nature of our data, we run our filters on each slice separately by only using 2 instead of 3 coordinates, by setting one of the two (U,V) coordinates to a constant value (e.g. V=1). For an extensive explanation on how the coordinates are computed, see Gulban et al., 2022. Although this approach for a volumetric parametrization of the cortex has been previously used in some recent layer-fMRI papers to investigate mesoscopic spatial features (Dresbach et al., 2024a, 2024b; Pizzuti et al., 2023, 2024), we extended it to microscopy data for the first time.

### 2.5 AHEAD laminar signal extraction

We selected one specimen from AHEAD dataset (ID: 122017) for the highest signal and reduced artifacts in the occipital part of the brain. This specimen had already been reconstructed and aligned with the respective qMRI data at nominal resolution of 0.15 × 0.15 × 0.2 mm by original authors (Alkemade et al., 2022). For the microscopy part of the dataset, we applied three intensity normalization steps in order to enhance fine scale details and remove residual acquisition artifacts within the cortical landscape in the 2D microscopy slices. First, we performed a slice-by-slice intensity normalization based on percentile computation: for each voxel, we subtract the 5th percentile and divide by the difference between 95th and 5th percentile (**Supplementary Figure 1**). In this way, the intensity range is uniform across slices and it is normalized between 0-1. Second, we ran a geodesic low-pass filter with cylindrical kernel (radius: and height: 10% of the local cortical depth measurement) by using `LN2_UVD_FILTER` -median within Laynii for removing high-frequency artifacts (e.g. tears, cracks), that we called ‘Tears Filter’. **Figure 3** shows three examples of how the filter mitigates the presence of the artifacts, which are irreversible distortions in the histology field that are induced by cutting, mounting and staining (Fischl and Sereno, 2018, Chapter 4). Finally, we estimated local field bias around each voxel by using the same LayNii program (`LN2_UVD_FILTER` -median) but with a cylindrical kernel with a larger volume (radius: 0.5 and height: 100% of local cortical depth) and mitigated its effect by dividing the voxel intensity by this estimated field. Lastly, we introduced a new filter ‘Cutting Angle Filter’ for automatically detecting pieces of the gray matter to exclude from our successive laminar analysis that didn’t align well with the coronal cutting place (**Figure 4**). This misalignment is due to the fact that the angle of sectioning relative to the local tissue orientation crucially affects the resulting shape of the lamination pattern. To compare various brain areas, cutting sections had to have a consistent orientation relative to the cortical surface. An angle of 90° is considered optimal to extract cross-section information. Already in the early twentieth century, Von Economo and Koskinas were aware of this problem and conducted their seminal staining work by dissecting each gyrus and sulcus perpendicularly to its axis (Fischl and Sereno, 2018, Chapter 2). Following the same rationale, we developed the ‘Cutting Angle Filter’ as an alternative algorithmic solution. For our filtering procedure, we computed the local tissue orientation by using `LN2_LAYERS` -streamlines program from LayNii. This program outputs a 4D NIFTI containing a vector map that attributes to each voxel a radial vector connecting inner and outer gray matter surfaces (locally orthogonal to the two surfaces). Since in this case the 3D geometrical nature of the cortex is used to compute the vector map, we input the segmentation computed on the 3D reconstructed model (**Figure 4, A**). Then, we obtained an angular map (scalar) (**Figure 4, D**) by computing the angle between each voxel’s local orientation (**Figure 4, B**) and the vector along the cutting place (coronal) (**Figure 4, C**). Finally, we excluded each voxel whose angle exceeds 150° (**Figure 4, E**). According to Schleicher and colleagues (Schleicher et al., 1999a), a deviation of a maximum 60° from the vertical (90°) was accepted to not alter the laminar pattern. Finally, we quantified the ability of each microscopy contrast (Bielschowsky, Thionin, Parvalbumin) to differentiate visual areas by calculating a measure of similarity as the pairwise correlation between each laminar profile (see **Table 1**). We computed a statistical inference (ANOVA one way) to assess if the correlations across ROIs are different across the three staining contrasts. This was followed by performing a pairwise comparisons post-hoc Wilcoxon signed-rank test. For the qMRI part, the main artifact for the 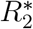 map within AHEAD dataset was the presence of vessel residual artifacts that appear as very bright “bubbles” in the data. This is due to the air remaining trapped in the vessels when preparing a post-mortem sample for imaging (Fischl and Sereno, 2018, Chapter 4). We used an intensity-based histogram matching algorithm from ITK-SNAP segmentation tools to detect and exclude affected voxels from further analysis.

**Table 1.** Similarity of laminar profiles (rho). Correlations are computed to quantify similarity across laminar profiles between each ROI (V1, V2, V3, hMT+) for each microscopy contrast. Low correlation values indicate high dissimilarity between laminar profiles of different regions. Significant differences are found between Parvalbumin and Bieloschowsky and Parvalbumin and Thionin and are highlighted by a red asterisk (p 0.01).

| (LH) | Bielschowsky | Thionin | Parvalbumin |
| --- | --- | --- | --- |
| V1-V2 | 0,994 | 0,994 | 0,997 |
| V1-V3 | 0,997 | 0,985 | 0,989 |
| V2-V3 | 0,996 | 0,996 | 0,991 |
| V1-hMT+ | 0,993 | 0,961 | 0,942 |
| V2-hMT+ | 0,995 | 0,963 | 0,921 |
| V3-hMT+ | 0,996 | 0,948 | 0,917 |
| (RH) | Bielschowsky | Thionin | Parvalbumin |
| V1-V2 | 0,999 | 0,999 | 0,995 |
| V1-V3 | 0,993 | 0,998 | 0,978 |
| V2-V3 | 0,996 | 0,999 | 0,974 |
| V1-hMT+ | 0,998 | 0,988 | 0,938 |
| V2-hMT+ | 0,998 | 0,989 | 0,925 |
| V3-hMT+ | 0,995 | 0,989 | 0,981 |

**Figure 3.**
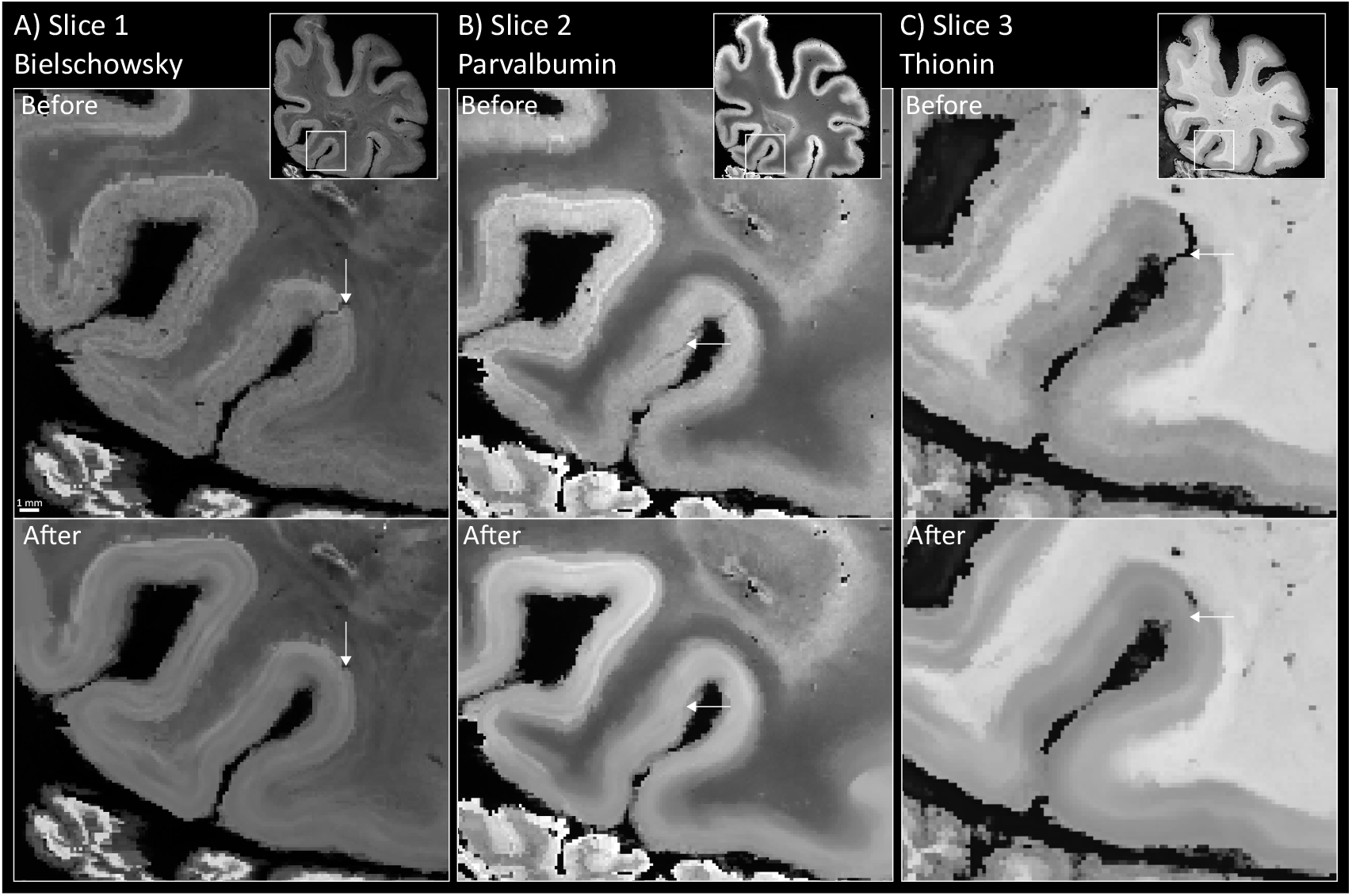
Tears filter. (A-B-C) Three exemplary slices spanning the three microscopy contrasts (Bielschowsky, Parvalbumin, Thionin) showing before and after the application of the tears filter. White arrows pointed to the tears that we aimed to remove. Square inserts showed the entire slices from which we zoomed in to highlight the artifact.

**Figure 4.**
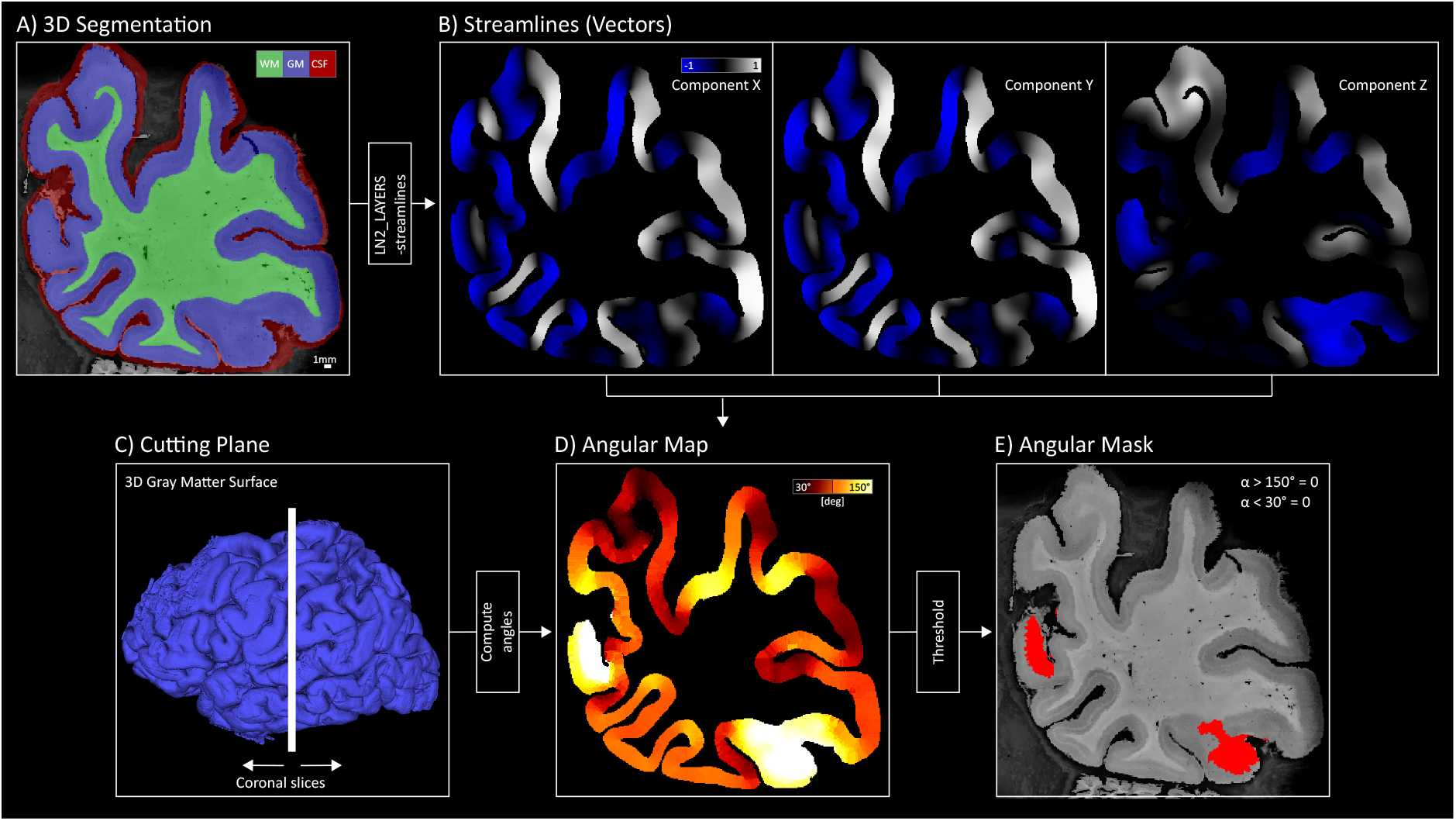
Cutting angle filter steps. A) The 3D segmentation file is used as input for computing the streamlines using the program `LN2_LAYERS` with -streamlines flag. A tissue segmentation overlaid on a Nissl stained slice is shown as an example. Colors indicate the three tissues, respectively: green for white matter, blue for gray matter and red for cerebrospinal fluid. Note that in post-mortem data the cerebrospinal fluid is physically not present and here indicates the outer boundary of gray matter. B) Streamlines vector field. The three components (x, y, z) of the vector field indicate the local tissue orientation for each voxel are shown for the exemplary slice. C) Schematic visualization of the cutting plane providing coronal slices as a vertical white line overlaid to the gray matter surface. D) The angular map represents the output of the angular computation between streamlines (B) and cutting plane (C). E) A threshold version of the map is shown as an angular mask; red patches indicate excluded areas for an angular measure exceeding 60° with respect to the cutting plane.

### 2.6 In-vivo MRI and fMRI laminar signal extraction

In order to preserve the fine scale details of our high-resolution 3D ME EPI anatomical images from processing resampling steps, we firstly upsampled each run to double resolution (0.175 iso. mm), as previously done by (Gulban, 2024). We used the -upsample function from the greedy package with nearest-neighbor interpolation (Yushkevich et al., 2006). For each run separately, we computed the average of the three echos and used them as reference to bring the runs to the same space by using the linear and the non linear registration program from greedy. The transformation matrix was then applied to each echo separately. Again, nearest neighbor was used as an interpolation step. Finally, we averaged the four runs and fitted a monoexponential decay function to compute the quantitative 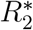 map. The anatomical images with UNI contrast at 0.35 iso. mm resolution, resulting from slab-stitching procedure, underwent the following processing steps: first, we applied a structure tensor denoising algorithm (Gulban et al., 2018) to increase the SNR (Gulban, 2024). Then, we upsampled to 0.175 iso. mm (as done for ME 3D EPI data) and we registered to the ME 3D EPI space using a non-linear co-registration procedure as implemented in greedy. Finally, we resampled our data using linear interpolation and used the resulting output as input to define cortical surfaces, ROIs and tissue segmentations. Resting-state fMRI (rs-fMRI) data at 0.8 iso. mm underwent the following preprocessing steps: slice time correction (BrainVoyager), motion correction (BrainVoyager), distortion correction (FSL TOPUP), high-pass filter with 3 cycles (BrainVoyager). We averaged the time series and upsampled to 0.175 iso. mm to match anatomical resolution using the ndimage.zoom command from scipy (Virtanen et al., 2020) with spline interpolation (order 3). We coregister rs-fMRI data to high-resolution 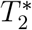 anatomical images (ME 3D EPI space) using a non-linear registration algorithm as implemented in greedy with linear interpolation.

### 2.7 Estimation of scaling factor between post-mortem and in vivo 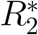 laminar profiles

To evaluate the relative contribution of microstructure and vasculature in the resulting 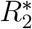, as the signal shifts from being influenced solely by microscopic factors (post-mortem) to a combination of microscopic and vascular contributions (in-vivo), we estimated a scaling factor between quantitative 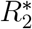 laminar profiles between the two datasets. While brain fixation has been shown to significantly affect the *R*_1_ range (Dinse et al., 2015), its impact on 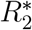 is considered minimal (Deistung et al., 2016). Given the similar 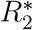 ranges observed, we attribute the differences between in vivo and post-mortem 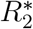 primarily to variations in tissue composition. For each laminar profile (four ROIs, two hemispheres, two datasets), we fit a linear regression model. For each ROI, we averaged the slope between the two hemispheres. Then, we divided the slope fitted on the post-mortem by the slope fitted on the in-vivo data. This ratio estimates the scaling factor of the 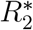 laminar profiles between the two modalities.

## 3 Results

### 3.1 Comparing the microscopic cortical architecture of visual areas

We report the lamination patterns of four visual areas, V1, V2, V3, and hMT+, using three microscopy contrasts: Bielschowsky, Thionin, and Parvalbumin (**Figure 5** and **Supplementary Figure 3**). Bielschowsky staining correlates with fiber density and indirectly with myelin, while Thionin and Parvalbumin provide cellular information by staining neuronal cell bodies and a subset of inhibitory interneurons, respectively. This analysis enables us to characterize each ROI by comparing the expression of these three microscopic features across cortical depths. We quantified the ability of each microscopy contrast to differentiate visual areas by calculating a measure of similarity (indexed as the pairwise correlation between each laminar profile (**Table 1**). Parvalbumin exhibited the lowest level of similarity across regions compared to Bielschowsky and Thionin, which was also qualitatively visible when comparing laminar profiles (**Figure 5** and **Supplementary Figure 3**). To support our comparisons, we tested if the correlations across the three groups are different: a one way ANOVA shows significant differences across groups (F = 9.03, p = 0.0007). A pairwise comparisons using post-hoc Wilcoxon signed-rank test revealed significant difference (after Bonferroni correction) between Parvalbumin and Bielschowsky (p = 0.002) and Parvalbumin and Thionin (p = 0.02) and not significant between Bielschowsky and Thionin (p = 0.1). Unlike cellular neuronal density, the distribution of Parvalbumin neurons varies characteristically across cortical depths and ROIs. The different distribution of parvalbumin neurons across layers may suggest a differential contribution to inhibitory processing with a maximum for intermediate layers.

**Figure 5.**
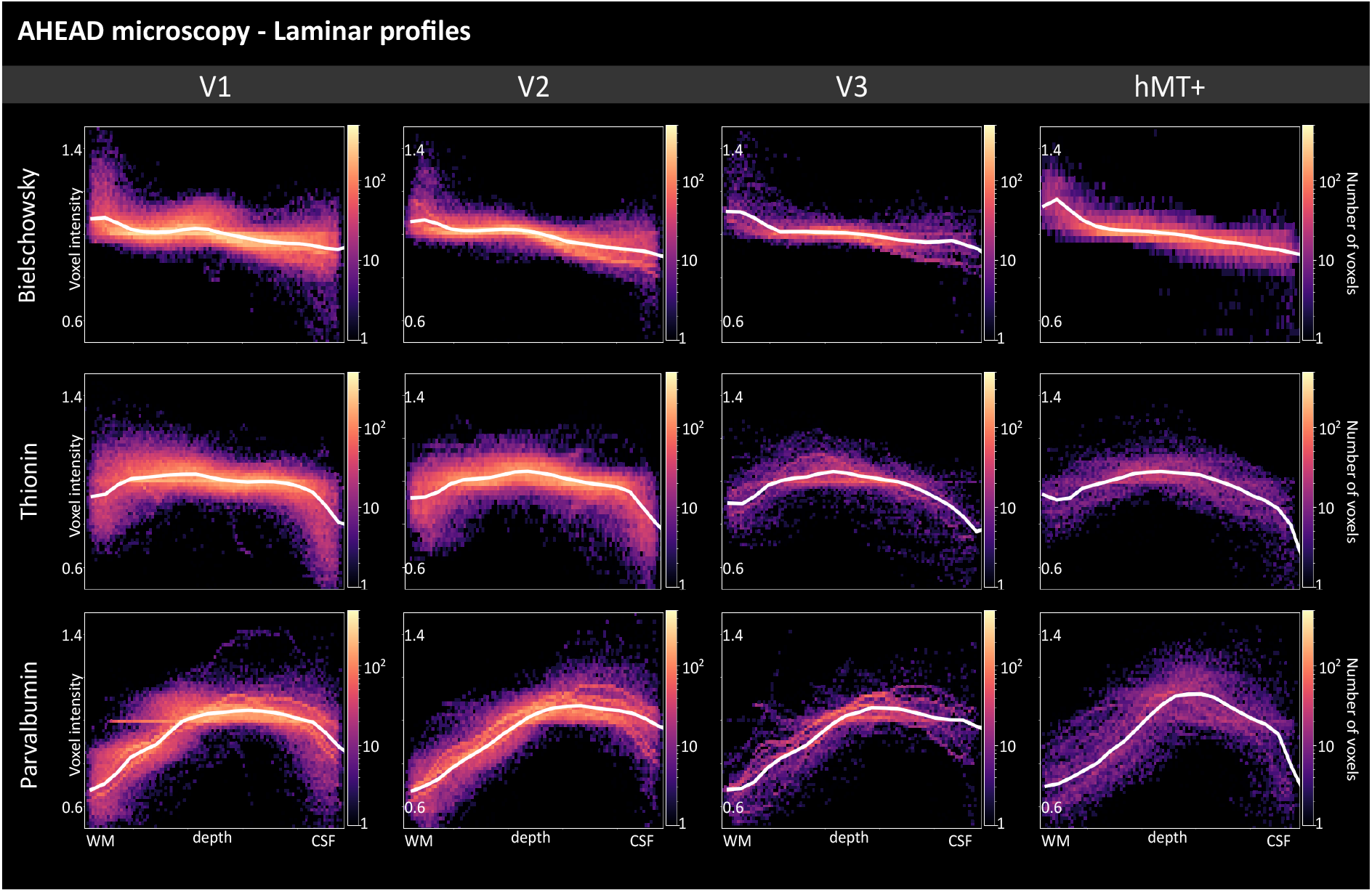
AHEAD laminar profiles for three microscopy contrasts (Bieloschowsky, Thionin, Parvalbumin) are shown as 2D histograms for each ROI of the left hemisphere. Gray matter cortical depth measure is shown from white matter (x=0) to cerebro-spinal fluid (x=1) boundary from left to right. Solid white lines in each subplot show median intensity for 21 discrete equivolume layers. The Y-axis is shown within the 0.5-1.4 (a.u.) range for each subplot. Results from the right hemisphere are shown in **Supplementary Figure 3**.

### 3.2 Comparing post-mortem and in-vivo quantitative 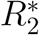 laminar profiles

Bridging the gap between post-mortem and in-vivo studies is crucial for understanding the sources of 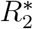,as it is assumed that the signal shifts from being influenced solely by microscopic factors (post-mortem) to a combination of microscopic and vascular contributions (in-vivo). Therefore, in this study, we report the quantitative 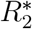 values across cortical depths for both post-mortem and in-vivo datasets (**Figure 6** and **Supplementary Figure 4**). **Table 2** accompanies **Figure 6** and **Supplementary Figure 4** by reporting the average 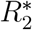 value in only three equivolume layers. First, our 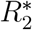 cortical layer profiles matches with the previously reported 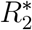 values within the visual cortex (Gulban et al., 2022). Second, we also observe overall matching profiles between the AHEAD and in-vivo data that shows a consistent decrease of 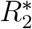 values from deep to the superficial layers. When comparing the magnitude of the 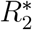 laminar profiles between post-mortem and in-vivo data, we show that the AHEAD 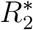 values are lower compared to those measured in vivo. In addition, the discrepancy between the 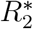 values seem to be the highest in superficial layers. This discrepancy is expected, as in-vivo 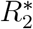 values include the additional contribution from blood vessels, leading to a faster signal decay. In contrast, post-mortem 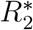 is influenced only by iron and myelin content, since the brain fixation process removes most of the blood, resulting in a slower decay. These differences are most pronounced from the middle to the superficial layers, which is expected due to the presence of more vasculature and partly by pial veins (partial volume effect). We estimate an average scaling factor across our visual ROIs between post-mortem and in-vivo of 2.8 (**Table 3**). This means that, the blood affects the estimation of 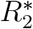 with a factor of three: the slope of the 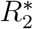 laminar profiles in vivo are almost three times smaller compared to the one measured when microstructure only is considered (post-mortem).

**Table 2.** Mean quantitative 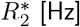 [Hz] reported for three equivolume layers (deep, middle, superficial) for all the regions of interest in both post-mortem and in-vivo data.

|  | **R2\* [Hz] Post-mortem** |  |  | **R2\* [Hz] In-vivo** |  |  |
| --- | --- | --- | --- | --- | --- | --- |
| **(LH)** | **Deep** | **Middle** | **Superficial** | **Deep** | **Middle** | **Superficial** |
| **hMT+** | 30,91 | 26,08 | 16,37 | 33,78 | 30,81 | 27,34 |
| **V1** | 30,83 | 26,25 | 18,12 | 33,97 | 32,11 | 29,14 |
| **V2** | 34,48 | 29,29 | 19,65 | 34,5 | 31,02 | 28,47 |
| **V3** | 33,87 | 29,21 | 18,94 | 32,96 | 29,74 | 28,05 |
| **(RH)** | **Deep** | **Middle** | **Superficial** | **Deep** | **Middle** | **Superficial** |
| **hMT+** | 32,62 | 27,11 | 17,27 | 34,23 | 31,3 | 26,93 |
| **V1** | 34,18 | 28,24 | 19,58 | 32,92 | 31,35 | 29,34 |
| **V2** | 37,82 | 31,56 | 20,41 | 34,38 | 31,18 | 28,55 |
| **V3** | 38,53 | 20,54 | 17,15 | 33,15 | 29,33 | 27,11 |

**Table 3.** Regression modeling for 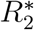 laminar profiles. The slope of the fitted curve is reported for each ROI. The ratio is considered as an estimate of the scaling factor between the two modalities.

| Post-mortem (slope) | V1 | V2 | V3 | hMT+ |
| --- | --- | --- | --- | --- |
| Left hemisphere | -22,39 | -24,87 | -21,2 | -24,74 |
| Right hemisphere | -24,01 | -28,51 | -34 | -26,33 |
| Average | -23,2 | -26,69 | -27,6 | -25,53 |
| **In-vivo (slope)** | **V1** | **V2** | **V3** | **hMT+** |
| Left hemisphere | -8,48 | -8,91 | -7,95 | -10,01 |
| Right hemisphere | -7,37 | -10,11 | -9,43 | -11,16 |
| Average | -7,92 | -9,51 | -8,9 | -10,55 |
| **Ratio: Post-mortem / In-vivo** | 3 | 2,8 | 3,1 | 2,4 |

**Figure 6.**
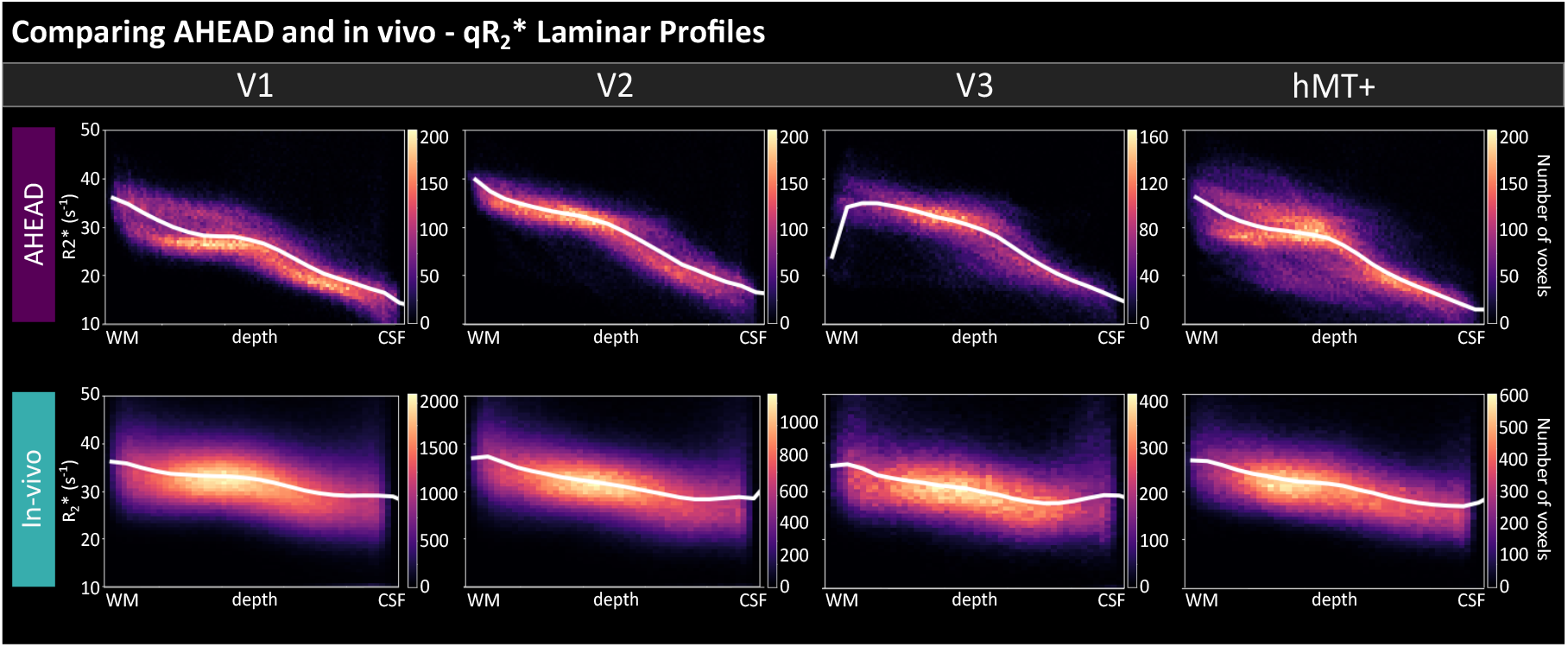
AHEAD (top) and in-vivo (bottom) 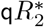 laminar profiles are shown as 2D histograms for each ROI for the left hemisphere. Gray matter cortical depth measure is shown from white matter (x=0) to cerebro-spinal fluid (x=1) boundary from left to right. Solid white lines in each subplot show median intensity for 21 discrete equivolume layers. The Y-axis is shown within the 10-50 (s-1) range for each subplot. Results from the right hemisphere are shown in **Supplementary Figure 4**.

### 3.3 Comparing structure to function: anatomical 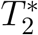 and resting state fMRI

Since GE-BOLD fMRI signals reflect variations of 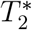 between conditions, we analyzed both anatomical and functional signal as 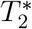 variation 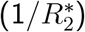.Here, our goal is to report the laminar profiles from the rs-fMRI run and to compare it to the anatomical 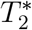 laminar profiles from both in-vivo and post-mortem AHEAD dataset (**Figure 7** and **Supplementary Figure 5**). In all ROIs, the rs-fMRI signal exhibits a general increase towards the cortical surface, similar to the anatomical profiles in the two datasets. Although at half of the spatial resolution (0.8 iso. mm), the same overall characteristic positive slope laminar feature we observed in the anatomical 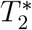 is also present within the rs-fMRI data. However, finer details observable in the anatomical laminar profiles, such as the characteristic dip in 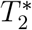,remain concealed in the current resting state laminar profiles due to the varying spatial scales. Future fMRI studies employing higher spatial resolution may uncover these details (Feinberg et al., 2023).

**Figure 7.**
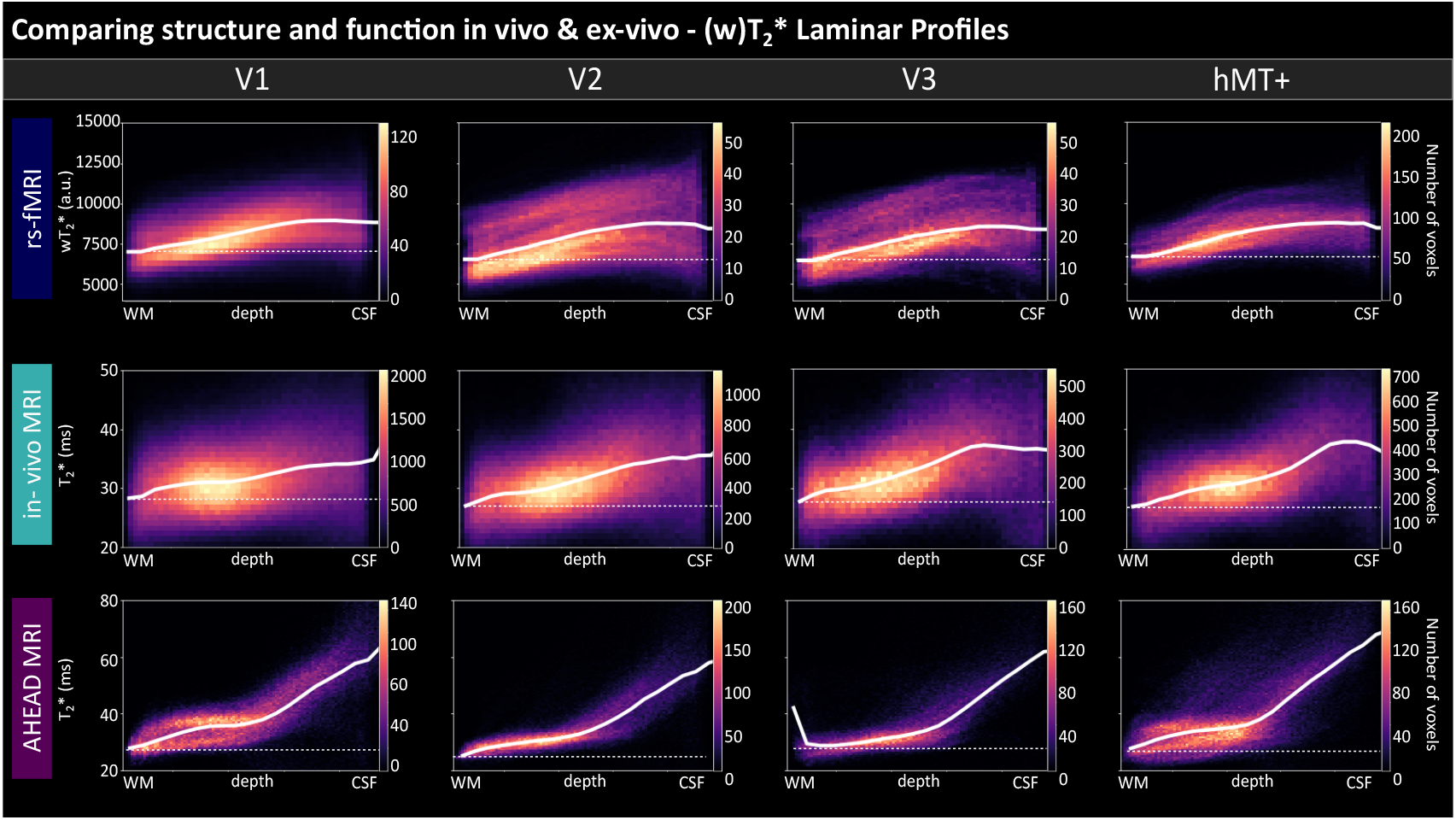
Structure-to-function comparison. Resting-state fMRI (top), in-vivo 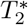 (middle) and post-mortem AHEAD 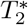 (bottom) laminar profiles are shown as 2D histograms for each ROI. Gray matter cortical depth measure is shown from white matter (x=0) to cerebro-spinal fluid (x=1) boundary from left to right. Solid white lines in each subplot show median intensity for 21 discrete equivolume layers. Dotted lines indicate horizontal lines to highlight the increase towards superficial layers of main curves. The Y-axis range is displayed only in the first subplot of each data type and kept invariant across ROIs. Results from the right hemisphere are shown in **Supplementary Figure 5**.

## 4 Discussion

### 4.1 Summary

In this study, we propose a multimodal laminar characterization for four human visual areas (V1, V2, V3, hMT+) that aims to bridge the micro and mesoscale. We used the novel publicly available post-mortem AHEAD dataset, which uniquely presents multiple microscopy contrasts and quantitative MRI for the same individual, and complement it with our in-vivo dataset consisting of both high-resolution anatomical and functional MRI (**Figure 1**). We investigated the microscopic underpinning of our regions of interests by analyzing the cortical variation of three microscopy contrasts (Bielschowsky, Thionin, Parvalbumin) and found a central role for Parvalbumin for area differentiation (**Figure 5, Supplementary Figure 3**). Moreover, we compared 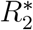 MRI cortical variation of post-mortem to in vivo samples, and found a common linear decrease 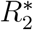 across the two modalities (**Figure 6, Supplementary Figure 4**). Finally, we report the laminar profiles in rs-fMRI (**Figure 7, Supplementary Figure 5**) and compare it with the structural laminar profiles measured in the same living brain and in post-mortem brain. Although at different spatial resolution, the same overall characteristic positive slope characterized the three laminar profiles.

### 4.2 Implications for layer-fMRI

#### Cortical distribution of Parvalbumin interneurons

It is generally assumed that the neural activity of excitatory neurons is reflected in BOLD fMRI response, since excitatory neurons constitute 80-90% of all cortical neurons (Meyer et al., 2011). However, it is known that the ratio between excitatory and inhibitory neurons varies across cortical depth and areas (Markram et al., 2004; Tremblay et al., 2016). Even though our microscopy data cannot quantify the ratio of excitatory and inhibitory neurons, we still found it notable that the cortical distribution of Parvalbumin neurons is a characteristic feature of the region of interest (**Figure 5**). Parvalbumin neurons, even if they represent only one category of interneurons, they are the most abundant GABAergic neurons in the cortex (Rudy et al., 2011) and are assumed to have indirect influence through the inhibition of pyramidal cells, resulting in vasoconstriction (Lee et al., 2021). Due to this property, neural mechanisms involving this category of interneurons can affect the hemodynamic response detected by fMRI (Moon et al., 2021). In particular, when submillimeter resolution is available for layer-fMRI studies, unveiling the cortical distribution of parvalbumin interneurons can be crucial to interpret the laminar results when an inhibitory circuit is hypothesized to be recruited for solving functional tasks. For instance, (Torres-Gomez et al., 2020) suggested that changes in the proportion of parvalbumin neurons in layers 2/3 cells may favor the emergence of activity encoding working memory in association areas in primate brains. We expect that integrating a proxy for microstructural neural information in humans will improve the interpretation of laminar functional response in resolving complex tasks.

#### Vascular vs microstructural gray matter composition

When reaching submillimeter spatial resolution in fMRI, signals from different cortical laminae can be disentangled. However, at this spatial scale also other mesoscopic details regarding the underlying vascular network have to be considered while discussing layer-fMRI results. In particular, the BOLD signal reflects variation of 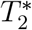 mainly coming from veins (Koopmans and Yacoub, 2019; Uludag and Havlicek, 2021). The presence of pial veins together with intracortical veins usually induces a bias known as ‘draining vein’ effect that is manifested as a signal that linearly increases towards the cortical surface. This is due to the fact that the blood is drained from deep to superficial layers by the veins that accumulate signals while traveling upwards, making the identification of the neural laminar source challenging (Koopmans and Yacoub, 2019). This cortical trend has been reported for both task-induced (Aitken et al., 2020; Fracasso et al., 2018; Mourik et al., 2021) and resting-state activity (Guidi et al., 2020; Markuerkiaga et al., 2021; Pais-Roldán et al., 2020). In this study, we showed that a linear trend characterizes the gray matter structure both in-vivo and post-mortem 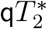 laminar profiles (**Figure 7**). When the vasculature is taken out of the equation, as in the post-mortem case, 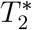 is still the highest at superficial layers. This result suggests that microstructural properties (e.g. myelin, iron) contribute to the linear increase in addition to vasculature in both task and resting state fMRI. Given that regional differences in the relative contributions of microstructure and vasculature are likely, understanding how these elements might consistently or differentially shape 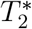 profiles across brain regions could enhance our interpretation of laminar fMRI. A scenario where both factors together produce a consistent linear increase across the cortex would align with prior observations, underscoring the importance of further investigation. Extending generative laminar models to embed area-specific microstructural and vascular features (such as 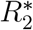 laminar profile) might complement and improve our understanding of the sources of variance driving layer-fMRI (Havlicek and Uludağ, 2020; Markuerkiaga et al., 2016).

#### Consequences for models of layer fMRI

Generative laminar models are useful tools to predict layer fMRI dynamics. However, the complexity and the accuracy of those models depends on the assumptions and on the physical parameters estimates (Havlicek and Uludağ, 2020; Markuerkiaga et al., 2016; Uludağ et al., 2009). Below, we discuss our resting-state laminar profiles together with the generative laminar model from (Havlicek and Uludağ, 2020). The model uses two compartments (intravascular (blood) and extravascular (parenchyma)) to describe the BOLD signal generated by a GE sequence (Eq. 1, Appendix). T1 effects are neglected in this model. As the signal from the intravascular compartment can be neglected at 7T (due to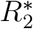), the predicted laminar profile depends only on the extravascular compartment. According to this modeling, the variables affecting the signal are:

1. S0 or rho, baseline signal intensity measured at TE=t(0), when no relaxation (particularly 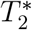 relaxation) has taken place. This signal measures the water proton density in the tissue.
2. CBV, cerebral blood volume, measuring the fraction of cerebral blood volume within a given amount of brain tissue.
3. 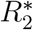,the transverse relaxation rate, which reflects how quickly the MRI signal decays in a gradient-echo sequence.

Conventionally, it is assumed that both S0 and 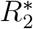 are constant across cortical layers and only CBV is expected to increase from deep to superficial layers (see Table 2, from Havlicek and Uludağ, 2020). Under this set of assumptions, the equation predicts a signal decreasing from deep to superficial layers (see Appendix - Laminar resting state fMRI). However, our empirical laminar resting-state fMRI profile shows the opposite trend (**Figure 7, Supplementary Figure 5**) with respect to the prediction. A similar trend to our results was also reported in previous works (Guidi et al., 2020; Markuerkiaga et al., 2016; Pais-Roldán et al., 2020). This discordance between modeling and empirical data points to the obvious conclusion that some of the above assumptions have to be relaxed in order to counterbalance the effect of CVB. In this work, we focus on quantitative 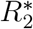 and consistently show in both post-mortem and in-vivo 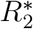 is clearly modulated (and not constant) with respect to cortical depth. The laminar variation of 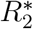 with its characteristic decrease from deep to superficial layers is one factor that counterbalances the effect of CBV: as 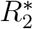 decreases with cortical depth, the 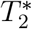 weighted rs-fMRI signal increases with cortical depth. Together with 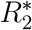,it is plausible to think that S0 also changes across cortical depth as a concurrent contribution to the resulting laminar profile measured during a rest condition. Empirical data, such as our observed laminar profiles, are invaluable in challenging and refining these models, offering insights that purely theoretical approaches may overlook. A full suite of modeling simulations is necessary to systematically explore the combined effects of both 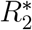 and S0 on the laminar profile. This approach can help resolve the observed divergence between current modeling predictions and empirical findings. By adjusting these parameters within the model, future work may better capture the nuanced interactions that shape the laminar BOLD signal across cortical depth, ultimately leading to more accurate representations of layer fMRI dynamics.

### 4.3 Extending open access tools for the AHEAD dataset

Publicly available post-mortem datasets are invaluable for studying the microscopic structure of the human brain (Alkemade et al., 2022; Amunts et al., 2013). However, developing specialized analysis toolboxes is also essential for correctly extracting information from these rare datasets and integrating it with other modalities. For instance, the Allen Brain Map portal (https://portal.brain-map.org/overview) is complemented by a suite of publicly available analysis tools (https://github.com/AllenInstitute). Following the same rationale, our streamlined analysis pipeline released as an open github repository (https://github.com/27-apizzuti/multimodal_layers.git), extends the open access analysis tools for the AHEAD dataset provided by Alkemade and colleagues, offering new tools for extracting laminar information. To enhance the laminar details and ensure robust cortical sampling from multiple slices, we developed the ‘cortical tears filter’ (**Figure 3**) and the ‘cutting angle filter’ (**Figure 4**). These tools are crucial for deriving reliable contrast- and ROI-specific laminar information. While we acknowledge the existence of other softwares addressing similar artifacts (Kindle et al., 2011; Mancini et al., 2020; Schleicher et al., 1999b), we highlight that our implementation is designed to encourage widespread use of AHEAD dataset specifically by offering tools that can be applied on the downloaded data. Note that the tissue segmentation step is not fully automated and requires expertise and manual work. This is a crucial step for accurate and precise layer profiles when working with very high resolution images. With all analysis methods and data used in this paper (histology, qMRI, and layer-fMRI) being publicly accessible, our framework offers a valuable resource for researchers conducting similar studies in other brain regions. By facilitating the study of cortical laminar structure and integrating this with functional information, our approach supports a deeper understanding of the structure-function relationships across the cortex, potentially uncovering unique regional patterns that contribute to brain function and disease.

### 4.4 Limitations and conclusions

Although our efforts to bridge scales and techniques have led to new insights and discussions on the laminar organization of four visual areas, it is worth pointing out the limitations of our results. Firstly, we focused on a limited subset of visual regions (V1, V2, V3, hMT+), which restricts the scope of our findings from drawing widespread conclusions on laminar features and visual hierarchy. Secondly, we defined the ROIs for V1, V2, and V3 using the visfatlas (Rosenke et al., 2021) with cortex-based alignment, and hMT+ was delineated based on macro-anatomical criteria outlined in (Huang et al., 2019). While we acknowledge that different methodologies can lead to partially overlapping regional boundaries (Turner, 2019 and Fischl and Sereno, 2018, Chapter 7), we argue that these discrepancies are unlikely to impact our results as we aim to understand regional and not local lamination differences. Additionally, comparing post-mortem and in vivo tissue introduces limitations due to the effects of brain extraction and fixation. These effects could influence the 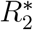 measurements (Deistung et al., 2016) and may alter the apparent thickness of cortical layers due to slight tissue shrinkage during fixation (Mouritzen Dam A., 1979). Finally, the small sample size, relying on data from only one post-mortem and one in vivo brain, limits the generalizability of our findings to broader populations, as individual anatomical variations may not be fully represented. However, the fundamental structural features observed in this study are likely to be consistent across human brains, supporting the broader relevance of our findings. Therefore, it is critical that similar post mortem and in vivo high resolution imaging efforts continue in the future where the amount of publicly available datasets increase. We conclude that our study represents a step forward along the goal of studying structure to function coupling mechanisms. Our results on regional laminar differences observed using multi-modal and multi-contrast data opens up new discussions on interpretation of layer-fMRI data. Future research could focus on expanding the multimodal characterization to include additional visual areas and functional contexts, enhancing our understanding of the dynamic interplay between micro- and mesoscale features in visual processing.

## 5 Appendix

### Laminar resting state fMRI

The laminar BOLD signal equation (Eq. 5 from Havlicek and Uludağ, 2020) for a cortical layer (denoted by subscript *k*) for a baseline condition (denoted by subscript 0) is given as:

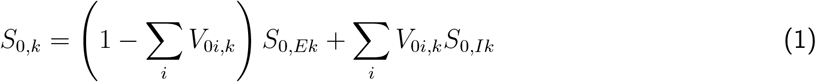

where:

1. *V*_0*i,k*_ is the cerebral blood volume (CBV) from the vascular compartment (denoted by subscript *i*) at baseline for the *k*-layer.
2. The vascular compartment include the venules and ascending veins compartments.
3. *S*_0,*Ik*_ is the intravascular signal at baseline for the *k*-layer.

### Laminar resting state fMRI

Under the approximation that, at 7T, 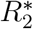 of the blood (intravascular) is twice as fast as 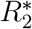 of the parenchyma, we can assume that *S*_rest0,*Ik*_ = 0 Havlicek and Uludağ, 2020.

Thus, we can write:

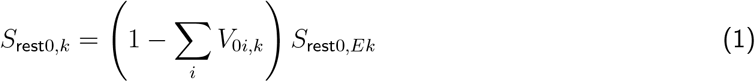

Since the parenchyma 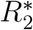 (extravascular) is assumed to be constant across cortical depth, we approximate *S*_rest0,*Ek*_ to be constant as well across cortical depth (Havlicek and Uludağ, 2020).

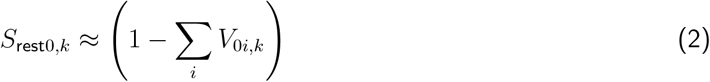

As CBV_0_ is expected to linearly increase across cortical depth (Havlicek and Uludağ, 2020), the resulting signal at resting state is expected to decrease as a function of cortical depth.

## 6 Data and Software availability statement

Analysis code is available on GitHub: https://github.com/27-apizzuti/multimodal_layers.git. Please refer to Alkemade et al., 2022 for raw post-mortem data. In-vivo anatomical data are shared in Zenodo: https://doi.org/10.5281/zenodo.14147820. Raw resting-state laminar-fMRI data are shared in Zenodo: https://doi.org/10.5281/zenodo.14164885. Preprocessed data from both post-mortem and in-vivo datasets are shared in Zenodo: https://doi.org/10.5281/zenodo.14164885.

## 7 Acknowledgements

This project was funded by the EU-project H2020-860563 euSNN and the European Union’s Horizon 2020 Framework Programme for Research and Innovation under the Specific Grant Agreement No. 945539 (Human Brain Project SGA3). In-vivo data was acquired at Scannexus (Maastricht, the Netherlands). OFG is funded by Brain Innovation. We thank Kamil Uludag for helpful discussions on the laminar modeling.SD is supported by the ‘Robin Hood’ fund of the Faculty of Psychology and Neuroscience and the department of Cognitive Neuroscience.

## 8 Author Contributions

According to the CRediT system (https://casrai.org/credit/)

**Conceptualization:** A.P., O.F.G., P.B.

**Methodology:** A.P., O.F.G., P.B.

**Software:** A.P., O.F.G.

**Validation:** A.P., O.F.G.

**Formal Analysis:** A.P.

**Investigation:** A.P., O.F.G.

**Resources:** A.P., O.F.G., P.B., J.P., R.G.

**Data curation:** A.P.

**Writing – original draft:** A.P.

**Writing – review & editing:** A.P., P.B., O.F.G., D.I., S.D., J.P., R.G.

**Visualization:** A.P.

**Supervision:** O.F.G., P.B.

**Project administration:** O.F.G., P.B., R.G.

**Funding acquisition:** R.G.

## 9 Supplementary Material

**Supplementary Figure 1:**
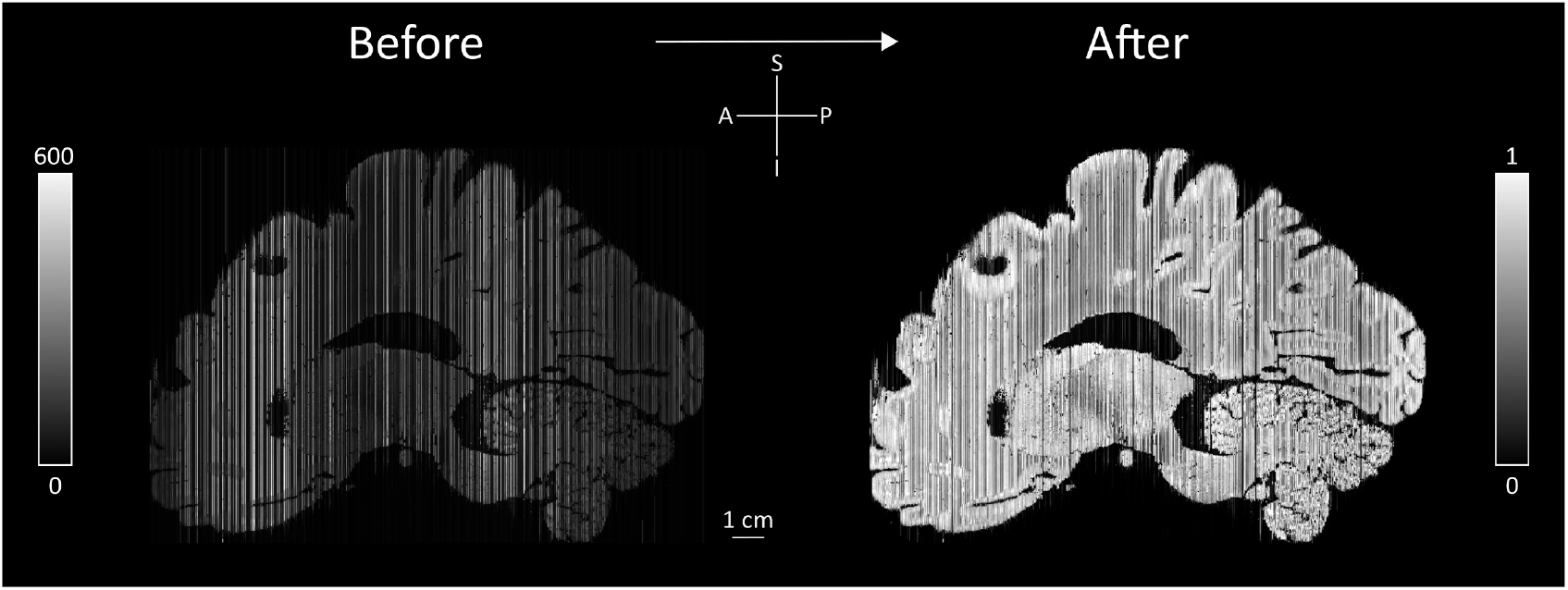
Slice-based percentile intensity normalization. Comparing AHEAD stack microscopy data (sagittal view) before and after this step. The intensity of each slice is normalized between 0-1.

**Supplementary Figure 2:**
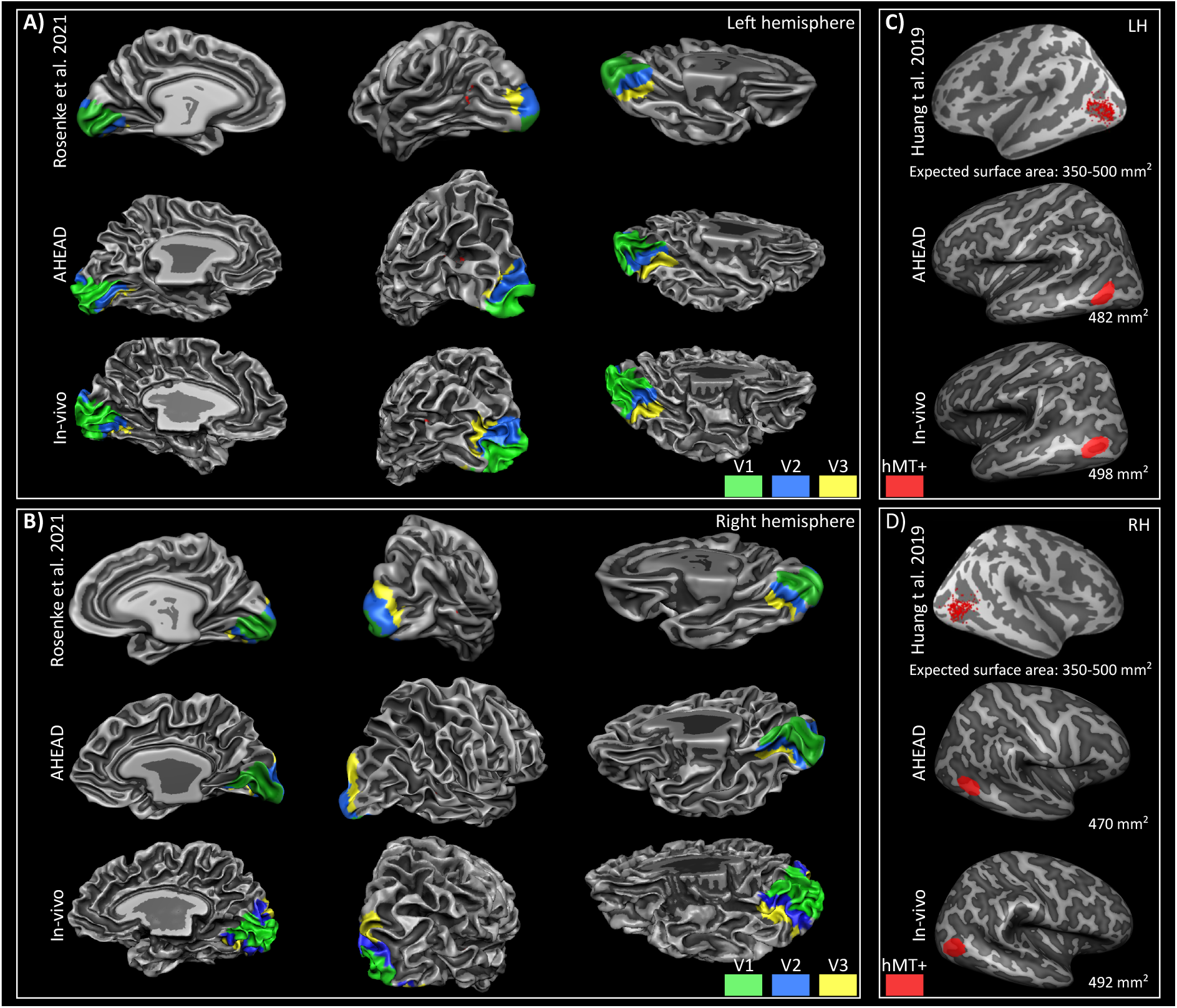
Extended ROI definition for both AHEAD and In-vivo datasets. Results from cortex-based alignment to the visual functional atlas (Rosenke et al. 2021) are shown on a white matter surface for both the left (A) and the right (B) hemisphere. Results from the macro-anatomical definition of hMT+ as explained in Huang et al. 2019 are shown on the inflated white matter surface for both the left (C) and the right (D) hemisphere. Surface area is reported for each hMT+ ROIs.

**Supplementary Figure 3:**
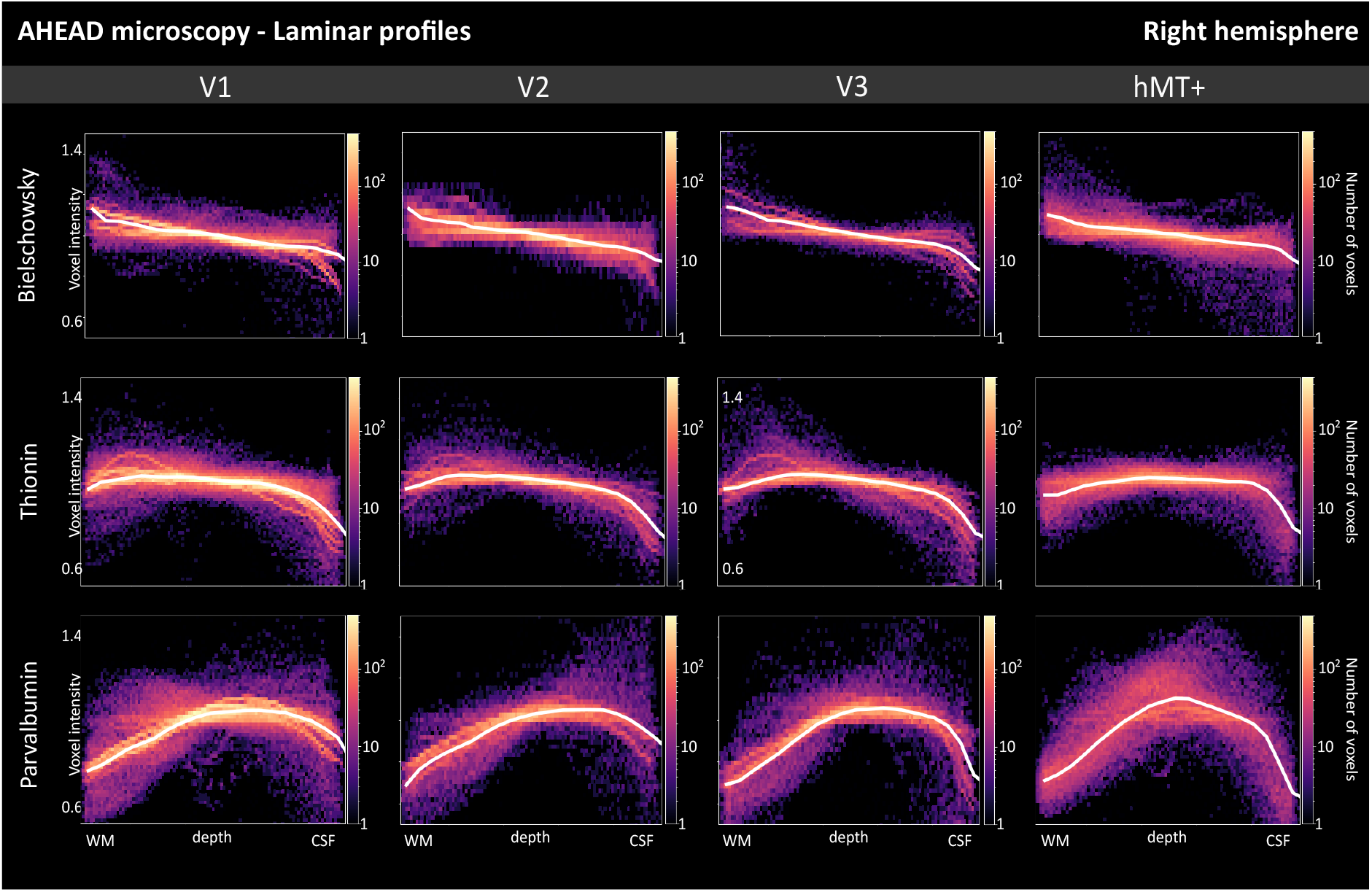
AHEAD laminar profiles for three microscopy contrasts (Bieloschowsky, Thionin, Parvalbumin) are shown as 2D histograms for each ROI of the right hemisphere. Gray matter cortical depth measure is shown from white matter (x=0) to cerebro-spinal fluid (x=1) boundary from left to right. Solid white lines in each subplot show median intensity for 21 discrete equivolume layers. The Y-axis is shown within the 0.5-1.4 (a.u.) range for each subplot. Results from the left hemisphere are shown in main Figure 5.

**Supplementary Figure 4:**
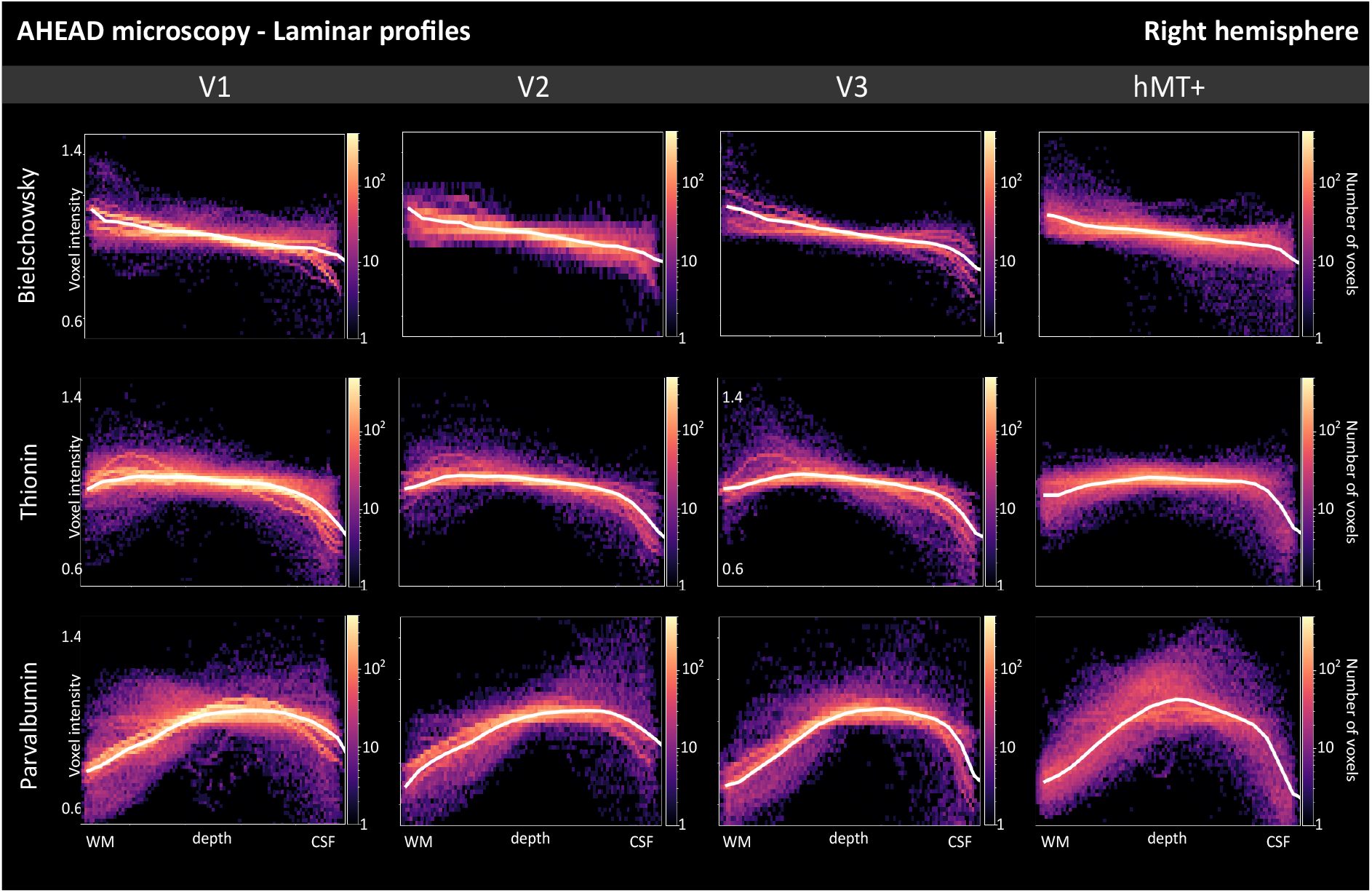
AHEAD (top) and in-vivo (bottom) 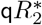 laminar profiles are shown as 2D histograms for each ROI for the right hemisphere. Gray matter cortical depth measure is shown from white matter (x=0) to cerebro-spinal fluid (x=1) boundary from left to right. Solid white lines in each subplot show median intensity for 21 discrete equivolume layers. The Y-axis is shown within the 10-50 (s-1) range for each subplot. Results from the left hemisphere are shown in main Figure 6.

**Supplementary Figure 5:**
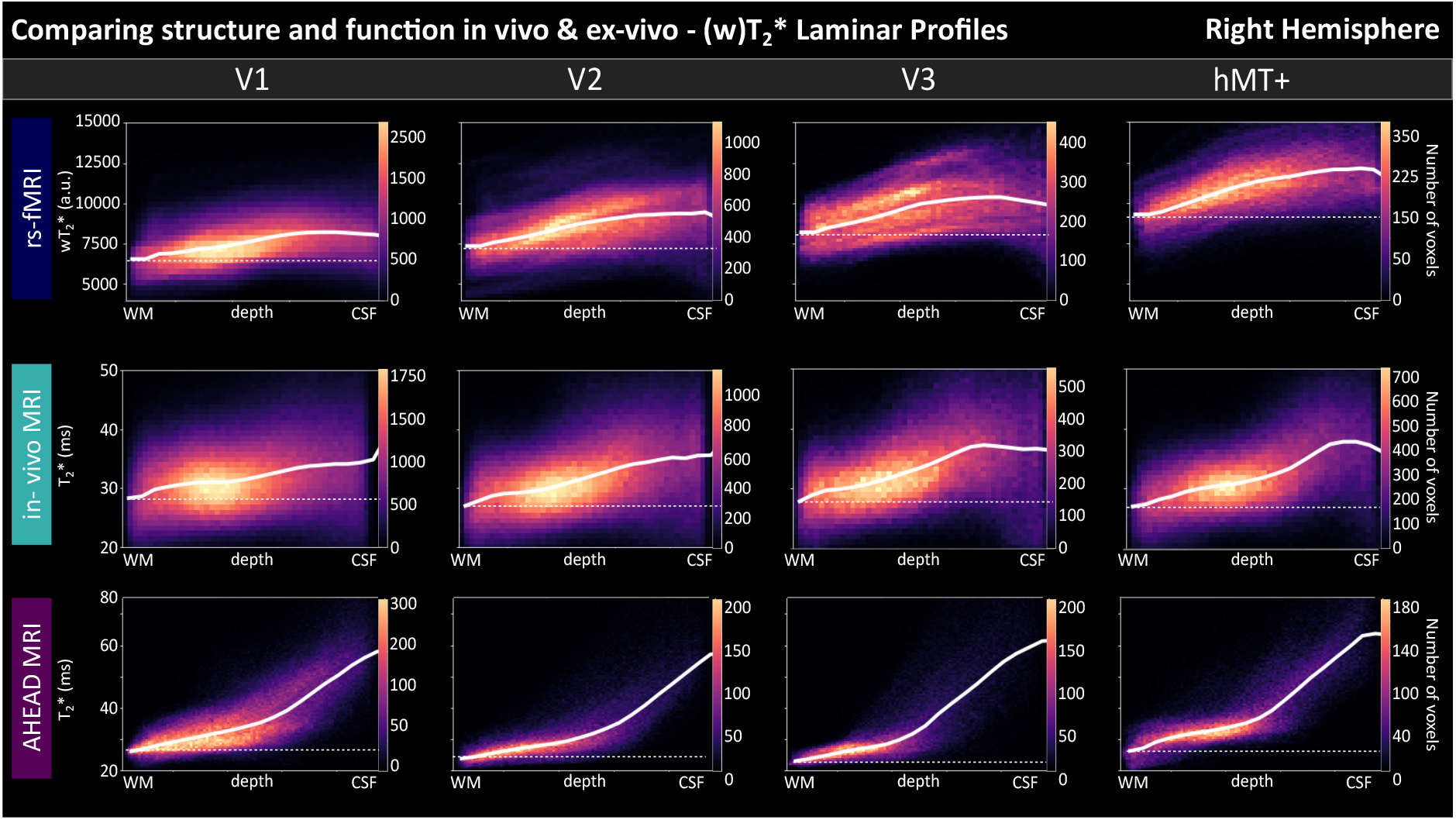
Structure-to-function comparison. Resting-state fMRI (top), in-vivo 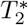 (middle) and post-mortem AHEAD 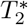 (bottom) laminar profiles are shown as 2D histograms for each ROI for the right hemisphere. Gray matter cortical depth measure is shown from white matter (x=0) to cerebro-spinal fluid (x=1) boundary from left to right. Solid white lines in each subplot show median intensity for 21 discrete equivolume layers. Dotted lines indicate horizontal lines to highlight the increase towards superficial layers of main curves. The Y-axis range is displayed only in the first subplot of each data type and kept invariant across ROIs. Results from the left hemisphere are shown in main Figure 7.

